# Rab converter DMon1 constitutes a novel node in the brain-gonad axis essential for female germline maturation

**DOI:** 10.1101/508598

**Authors:** Neena Dhiman, Girish Deshpande, Girish S Ratnaparkhi, Anuradha Ratnaparkhi

## Abstract

Monensin-sensitive 1 (Mon1) is an endocytic regulator that participates in the conversion of Rab5 positive early endosomes to Rab7 positive late endosomes. In *Drosophila*, loss of *mon1* (*Dmon1*) leads to sterility. The *Dmon1* mutant females have extremely small ovaries with complete absence of late stage egg chambers-a phenotype reminiscent of mutations in the insulin pathway genes. Consistently, we find that expression of many *Drosophila* insulin-like peptides (*Dilps*) is down regulated in *Dmon1* mutants. Conversely, feeding an insulin-rich diet can rescue the ovarian defects induced by the loss of *Dmon1*.

Surprisingly however, *Dmon1* is required in the tyramine/octopaminergic neurons (OPNs) and not in the ovaries or the insulin producing cells (IPCs). Thus, knockdown of *Dmon1* in just the OPNs is sufficient to mimic the ovarian phenotype while expression of *Dmon1* in the OPNs alone, is sufficient to ‘rescue’ the mutant defect.

Lastly, we have identified *dilp5* as a critical target of *Dmon1*. Both, protein and mRNA levels of *Dilp5* levels are reduced in *Dmon1* mutants and IPC-specific *dilp5* over expression can ameliorate the *Dmon1* dependent sterility defect. The study thus identifies *Dmon1* as a novel molecular player in the brain-gonad axis and underscores the significance of inter-organ systemic communication during development.

**Significance Statement:** Functional significance of the long-distance systemic communication during organogenesis has emerged as a major area of enquiry. We have focused our attention on an endocytic regulator DMon1, that appears to participate in a ‘remote control’ type of mechanism. We report a novel tripartite circuitry that involves *Dmon1* activity in Octopaminergic neurons, its influence on insulin production in the insulin producing cells (IPCs) which, in turn, is required for the progression of oogenesis. Our results document a spatially remote non-autonomous control mechanism involving neuronal cross-talk that orchestrates developmental regulation of oogenesis. Importantly our data provide a unique example of how distinct neuronal hubs can engineer metabolic pathways underlying growth/differentiation and highlight the importance of systemic regulation of organogenesis.

## Introduction

*Drosophila* oogenesis has served as an attractive and genetically tractable developmental model system with numerous distinguishing features (1–4). The two prominent traits that make it simple yet unique include exquisitely detailed patterning orchestrated by the dialogue between the germline and the surrounding soma, and fine-tuned coordination between the non-autonomous and autonomous factors that determine morphogenesis and growth (3, 5, 6).

*Drosophila* adult ovary is made up of 16-20 tubular structures termed as ovarioles (3, 7, 8). Each ovariole is comprised of six to eight egg chambers that develop in a sequential manner. Germarium at the anterior tip of each ovariole contains 2-3 germline stem cells (GSC) surrounded by terminal filament cells and cap cells that constitute the ‘niche’ i.e. the microenvironment necessary for both the self-renewal and, subsequent differentiation of the GSCs. Each GSC typically undergoes asymmetric division producing one germline stem cell daughter and a cytoblast. The newly formed cystoblasts undergo four incomplete mitotic divisions to form 16 interconnected cyst cells. Out of the 16 cells, one develops into the future oocyte, and the remaining 15 form the nurse cells. The future oocyte can be distinguished from the other cells based on early enrichment of transcripts and proteins such as *orb, BicD* and *Egl*.

Progression of egg chamber development is divided into 14 defined stages of maturation. All the egg chambers are connected to one another via stalk cells which are somatic in origin. Till stage 8, the size of the individual nurse cells is comparable to that of an oocyte. By stage 9 however, the oocyte begins to increase in size significantly. By the end of stage 14, as the egg chamber proceeds towards the posterior end of the ovariole, a mature oocyte passes through the lateral oviduct across the common oviduct and ultimately exits via the common oviduct (9).

The growth of the egg chamber thus can be broadly divided into the previtellogenic (*i.e*. upto stage 8) and vitellogenic phases (i.e. beyond stage 8). The vitellogenic phase is distinguished by the exponential growth of the oocyte due to yolk accumulation. The transition from pre-vitellogenic to vitellogenic stage is under tight hormonal as well as nutrient control. For instance, the Juvenile Hormone (JH) is required for yolk formation and necessary for vitellogenesis. This phase also involves extensive endocytosis by the oocyte (10). In this regard, it is interesting to note that vitellogenesis depends upon the coordinate control of growth by the fly’s nutritional status and signaling by insulin-like peptides. Interestingly, out of the 8 *Drosophila* insulin-like peptides (DILPs) three members namely DILP 2, 3 and 5 are produced by neurons termed as insulin producing cells or IPCs. These insulin-like peptides are released into circulation through the axon terminals of the IPCs at the corpora cardiaca-a neurohemal organ (11–13). Supporting the conclusion that DILPs exert a non-autonomous influence on the progression of oogenesis, germline specific loss of Insulin receptor (INR) or Chico (the insulin substrate protein) leads to a reduction in the size of the ovary and inability of the egg chambers to enter the vitellogenic stages and sterility (14–16).

Previous studies in *Drosophila* have indicated that IPCs are likely to be under neuronal regulation: Short neuropeptide F (sNPF) and octopamine appear to stimulate IPCs and thus can potentiate insulin signaling while GABA has inhibitory influence (17-19). Despite the fact that Insulin signaling affects a wide variety of developmental processes in several organismal contexts, mechanisms underlying insulin production, release and transport are not fully understood. The modes of short as well as long range transmission have been a focus of enquiry due to systemic influence of insulin signaling on growth and metabolism. It is generally believed that different regulatory circuits, both upstream and downstream, of insulin signaling must deploy a unique and dedicated set of regulators to achieve tissue specific outcomes. The molecular circuitry that participates during the synthesis and long-distance transmission of insulin to engineer proper *Drosophila* egg chamber growth and patterning are yet to be elucidated.

In this regard, we turned our attention to a highly conserved endocytic protein, Monensin sensitive 1 (hereafter referred to as Mon1). A protein complex between Mon1, a ‘longin domain’ containing protein, and CCZ1 functions as a guanine nucleotide exchange factor (GEF) for Rab7 (20). In a canonical endocytic cycle, Mon1 appears to be essential for the conversion of Rab5 positive early endosomes to Rab7 positive late endosomes (20, 21). Consequently, loss of Mon1 leads to accumulation and enlargement of early Rab5 positive endocytic compartment. In *Drosophila*, the process of Rab conversion is conserved and *DMon1* is involved in the recruitment of Rab7 (22). Intriguingly, at the neuromuscular junctions neuronal *Dmon1* is involved in regulating glutamate receptor levels on the post-synaptic side (23).

While characterizing P-element mediated excision mutations in *Dmon1* we noticed that homozygous mutant animals die throughout development while the escapers are short lived with severe motor defects (23). Recently, we also observed that *Dmon1* mutants display sex-non-specific sterility. Expectedly, the lethality and motor defects could be rescued by pan-neuronal expression of *Dmon1*. Curiously however, the neuronal expression is also capable of rescuing the sterility in these mutants suggesting that the neuronal expression of *Dmon1* may regulate fertility in wild type females.

Here, we have analyzed mechanistic underpinnings of the influence of *Dmon1* on oogenesis. Our findings have uncovered an unanticipated link between neuronal *Dmon1* activity and insulin production/signaling from the IPCs that has functional implications for the brain-gonad axis.

## Results

### Dmon1 mutations influence body size and ovarian length

We have previously reported generation of new alleles of *Dmon1* using P-element excision (23). Molecular characterization of one of these alleles *Dmon1^Δ181^* revealed that the C-terminal region from amino acid 249 onwards of *Dmon1* is deleted. Homozygous *Dmon1^Δ181^* mutants die throughout development and display motor defects (23). Moreover, *Dmon1^Δ181^* mutant adults are ~15% smaller in size (Fig.1A, B) compared to wild type adults of similar age and display sex-non-specific sterility. *Dmon1^Δ181^* females are 2188±97.07μm (n=9) compared to wildtype females which are 2535.82 ±72.27 μm (n=7) (Fig. 1A) in length. Sterility is fully penetrant in females and mutant females lay very few eggs, if at all.

**Figure 1.**
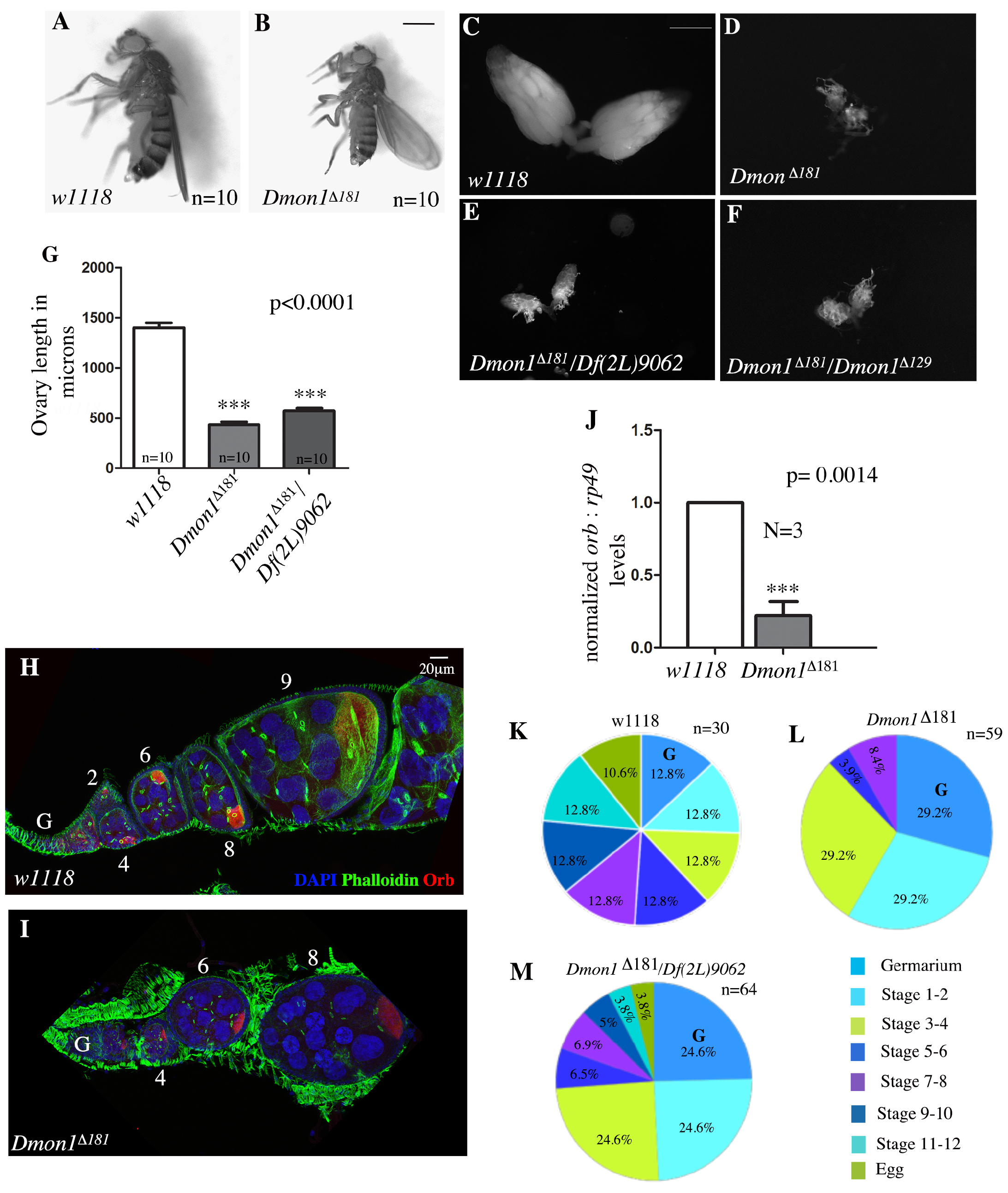
*Dmon1* mutants ovaries are smaller in size and are stalled at pre-vitellogenic stages. **(A, B)** A 2-3 day old wildtype (A) and age matched *Dmon1^Δ181^* animal. The mutant is ~22% reduced in size compared to wild type. **(C-G)** Ovary size is reduced by ~70% in *Dmon1* mutants. Ovary from a 2-3day old adult females of wildtype (C), homozygous *Dmon1^Δ181^* (D), *Dmon^Δ181^/Df(2L)9062* (E) *and Dmon^Δ181^/ Dmon^Δ129^* (F). Panel (G) shows the decrease in length of the ovary. p<0.0001, One-way Anova. **(H-I)** Wild type (I) and homozygous *Dmon^Δ181^* (J) ovarioles stained with orb (red), DAPI (blue) and phalloidin (green). Late stage egg chambers (stage 9 and 10) are absent in *Dmon1^Δ181^* mutants. **(J)** *orb* mRNA level, measured using real-time PCR is significantly lowered (approximately 80%) in *Dmon^Δ181^* mutants. **(K-M)** Pie-charts displaying the proportion of different stages of egg chambers in 2-3 day old *w1118*, homozygous *Dmon^Δ181^, Dmon^Δ181^/Df(2L)9062* adult females. Note the predominance of early pre-vitellogenic egg chambers in the mutants.

Upon dissection of the ovaries from 2-3-day old *Dmon1^Δ181^* mutant females along with control *w^Δ118^* females, it was immediately apparent that the mutant ovaries are remarkably small as opposed to the control (Fig. 1D). To exclude the possibility that this phenotype is genetic background dependent, we also examined ovary size in *Dmon1^Δ181^/Df(2L)9062* and *Dmon1^Δ181^/ Dmon1^Δ129^ females. Dmon1^Δ129^* carries a small deletion that spans the C-terminal region of *Dmon1* and the 5’ end of the neighboring *smog* (23). In both cases, the ovaries were found to be very small and comparable in size to *Dmon1^Δ181^* mutants (Fig. 1E, F).

Estimation of the length of the wild type and mutant ovaries from the proximal to distal end revealed that *Dmon1^Δ181^* ovaries (431.02±27.6 μm; n=10) were approximately 3 times smaller than the wildtype control (1400± 49.9μm, n=10; Fig.1G). A comparable decrease in size was also observed in *Dmon1^Δ181^/Df(2L)9062* adults (571.5±26 μm, n=10; Fig. 1G). Furthermore, the total number of ovarioles per animal was also significantly reduced in the *Dmon1^Δ181^*. In wild type the average number of ovarioles per animal is 35.05±0.4 (n=10), In contrast, the ovariole number in *Dmon1^Δ181^* mutants was *21.3 ±1.13* (n=5) corresponding to a decrease of approximately 35 %.

### Dmon1 mutant ovaries exhibit arrested egg chamber development and partial degeneration

To assess if different ovarian cell types (somatic as well as germline) are specified in the 2-3 day old mutant ovaries we stained *Dmon1* mutant ovaries with DAPI; a DNA dye and fluorescently tagged phalloidin to mark actin and actin-rich structures such as ring canals. We also co-immunostained these samples with antibodies against an oocyte marker, Orb. Ovaries from age-matched wild type animals were used as controls.

Expectedly individual *Dmon1^Δ181^* mutant ovarioles (Fig. 1I) are considerably smaller compared to the wild type (Fig. 1H). The level of Orb protein also appeared to be significantly diminished compared to the control. A quantitatively significant decrease in *orb* mRNA levels was also observed in *Dmon1* ovaries (approximately 80%; Normalized transcript abundance of 1 in wildtype versus 0.22±0.1 in *Dmon1^Δ181^ mutants* Fig. 1J).

The most striking phenotype seen in *Dmon1^Δ181^* mutants was the absence of late stage egg chambers. Most ovarioles had egg chambers of the pre-vitellogenic stages, with terminal egg chamber being of stage 7 or 8 (Fig. 1L). 10% of the mutant ovarioles also showed degenerating terminal egg chambers (data not shown). The developmental arrest of egg chambers was quantified by scoring each mutant ovariole for all the different stages. In a wild type ovariole, all the developmental stages starting with the germarium, till the mature egg are represented in roughly equal proportions (Fig. 1K). In contrast, in Dmon1^Δ181^ mutant ovarioles, arrest was observed from stage 5 onwards with a near complete absence of egg chambers beyond stage 8 (Fig. 1L, M). Consistently, a very low proportion of ovarioles displayed mature egg formation (Fig. 1L-M). Taken together these data suggest that the stalling defect observed in *Dmon1^Δ181^* mutant ovaries is a result of developmental arrest.

### Ovary specific knock down of DMon1 is insufficient to recapitulate the ovarian phenotype of Dmon1

To better understand the ovarian growth arrest phenotype induced by loss of *Dmon1*, we sought to determine the specific tissue and/or the cell type wherein *Dmon1* function is required using RNAi based reduction of *Dmon1* transcripts. We and others have previously used this approach successfully to compromise *Dmon1* function in a tissue/cell type specific manner (22, 23). To begin with, we employed *nanos (nos)-Gal4* and hearŕless(htl)^GĬMR93HỖ7^-Gal4 (24) to compromise *Dmon1* in the germ line and somatic epithelial sheath cells that surround the ovariole respectively (Fig. 2 A-D). The aspect ratio of the ovary i.e. length of the long axis divided by length of the short axis, was used as an indicator of size (greater the aspect ratio, smaller the size). Surprisingly, no appreciable change in size was observed upon driving RNAi using the germ line specific nos-GAL4: 1.52±0.04 (nos-GAL4/+; n=12) *vs* 1.53±0.05 (*nos-GAL4>UAS-Dmon1RNAi;* n=12) (Fig. 2E). Similarly, sheath cell specific knockdown of *Dmon1* using *htl* ^GMR93HỪ7^*-Gal4* was equally ineffectual. (htl-GAL4/+: 1.31 ±0.03, n=10 *vs htl-GAL4>UAS-Dmon1RNAi*: 1.41±0.06, n=10) (Fig. 2F). Further supporting the conclusion that ovarian tissue is unlikely to be the site of DMon1 action, knockdown of *Dmon1* using other ovary-specific drivers like *E22C-Gal4* (expressed in both the follicle cells and germline) *or maternal alpha-tubulin-GAL4:VP16* did not affect ovary size.

**Figure 2.**
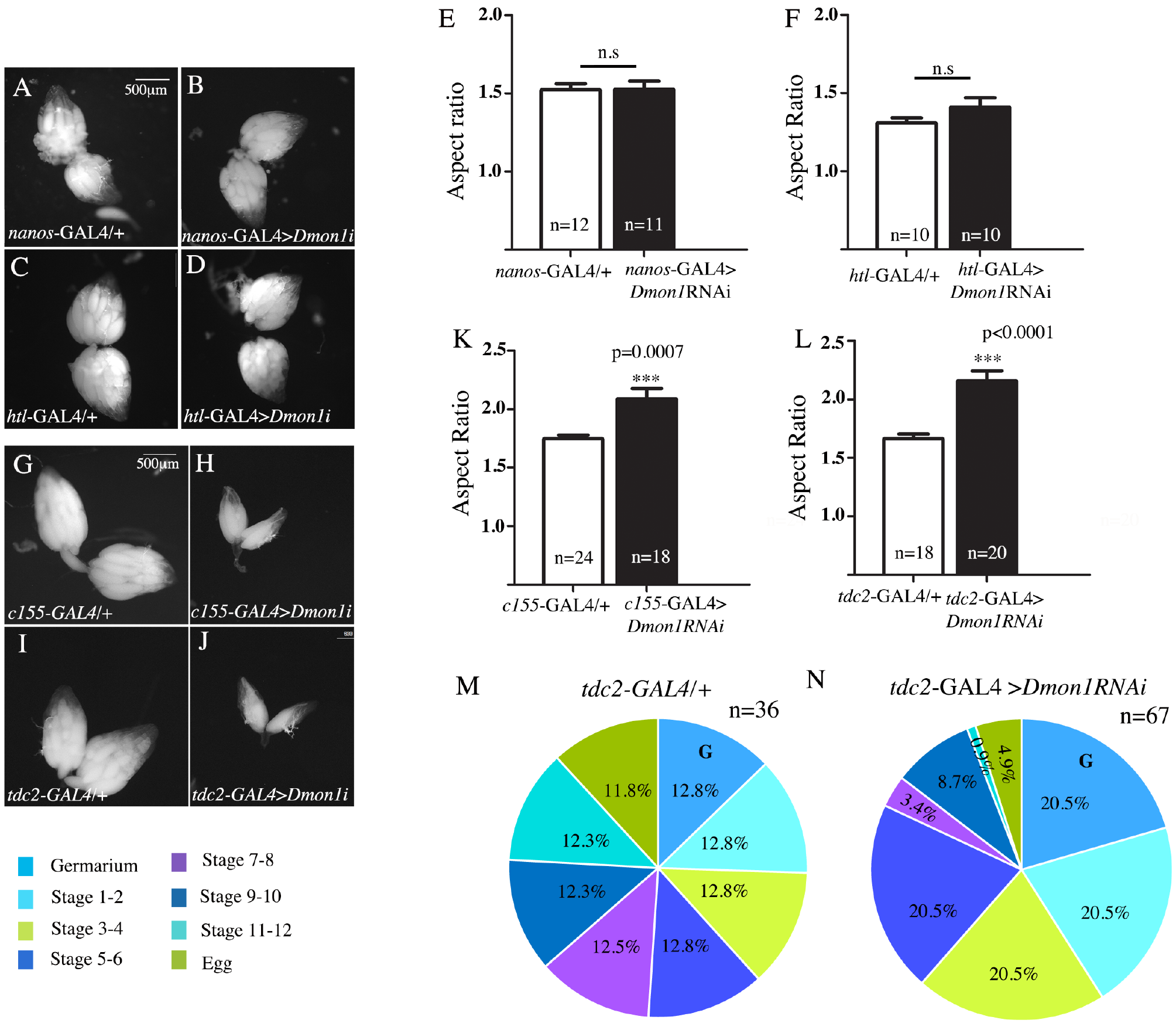
Knockdown of *Dmon1* in OPN neurons results in small ovaries with fewer late stage egg chambers. **(A-D)** Reduction of *Dmon1* transcripts in the germline (*nos*-GAL4>*UAS-Dmon1RNAi*) or soma (*htl*^GMR93H07^-GAL4> *UAS-Dmon1RNAi*) does not reduce ovarian size or perturb morphology. **(E, F)** Shown is the quantitation of the aspect ratio of the ovaries which is unaffected by either germline (E) or somatic (F) expression of *UAS-Dmon1RNAi*. **(G-J)** Pan neuronal (H) and *tdc2*-GAL4 specific knockdown of *Dmon1* (J) results in smaller ovaries compared to their respective controls (G and I). **(K, L)** Ovarian aspect ratios are significantly affected by neuronal knockdown of *Dmon1* transcripts in the *C155* and *tdc2* domains. **(M, N)** Pie-charts showing the distribution of stages of egg chambers in *tdc2*-GAL4/+ (M) and *tdc2*-GAL4>*UAS-Dmon1RNAi* (N) ovaries. Late stage egg chambers are underrepresented when *Dmon1* levels are reduced in the *tdc2*-GAL4 expressing neurons.

### Pan-neuronal inactivation of Dmon1 can partially mimic the ovarian phenotype of Dmon1 mutants

We have previously reported that pan-neuronal expression of *Dmon1* was able to alleviate lethality as well as the neuromuscular junction phenotype induced by loss of *DMon1* (23). We thus wondered if neuronal loss of *Dmon1* is sufficient to recapitulate some aspects of the ovarian phenotype of *Dmon1^Δ181^*. To test this idea, we expressed *Dmon1-RNAi* in the nervous system using a pan-neuronal driver, *C155-GAL4*. To our surprise neuronal expression resulted in significant reduction in the ovary size (Fig. 2H). The aspect ratio of these ovaries (Fig. 2K) was increased substantially, 2.09±0.09 (*c155-* GAL4/+> *UAS-Dmon1RNAi; n=18*) as compared to the ‘driver alone’ control 1.75±0.03 (C155-GAL4/+; n=18).

### Octopaminergic neurons are the likely site of *DMon1* action

Since *c155-GAL4* is a pan-neuronal driver, we wanted to determine if *Dmon1* is required in a subset of neurons. The appropriate candidate in this regard are the octopamiergic/tyraminergic neurons (OPNs) as the ovarian peritoneal sheath muscles are innervated by this class of neurons and there is no direct innervation to the ovary (25–27). We thus tested if knock down of *Dmon1* in OPNs alone, can influence ovarian growth and maturation. Indeed, knock down of *Dmon1* using *tdc2*-GAL4, which expresses in adult OPNs (28, 29), resulted in smaller ovaries (Fig. 2I-J) with an aspect ratio of 2.16±0.09 (tdc-GAL4> *Dmon1RNAi;* n=20) versus 1.67±0.04 (tdc-GAL4/+; n=18) (Fig. 2L). Given the decrease in size of the ovaries, we compared the distribution of egg chambers in *tdc2*-GAL4>*Dmon1RNAi* to ‘driver alone’ controls. Interestingly, a significant increase in the proportion of pre-vitellogenic egg chambers was observed along with a noticeable reduction in the number of mature eggs (Fig. 2M, N). Thus, knock down of *Dmon1* in OPNs is sufficient to partially mimic important aspects of the *Dmon1* mutant phenotype. Thus, our data suggest that *Dmon1* function in OPNs, is required for escaping developmental arrest. It should be noted that the RNAi dependent inactivation of *Dmon1* results in partially penetrant phenotype, possibly due to inefficient knockdown of the *Dmon1* transcript. Alternatively, it is possible that *Dmon1* function is also required in tissues other than the nervous system to influence oogenesis.

### Expression of *Dmon1* in the OPNs is sufficient to rescue the ovarian defects of *Dmon1^Δ181^* mutants

Pan-neuronal inactivation of *Dmon1* mimics the ovarian phenotype of *Dmon1*. Furthermore, pan-neuronal expression of *Dmon1* rescues lethality and sterility of *Dmon1^Δ181^* mutants (23). We therefore examined ovaries in *Dmon1^Δ181^* mutants rescued by expressing *Dmon1* using the pan-neuronal *c155-Gal4*. The ovaries from these animals looked normal in size and structurally similar to that in wild type animals (Compare Fig. 3B and Fig. 1C). Furthermore, the egg chamber arrest or ‘stalling’ phenotype observed in the mutants was completely rescued (data not shown). The ovaries from C155; *Dmon1^Δ181^/ Dmon1^Δ181^; UAS-Dmon1:HA* animals showed presence of egg chambers at all stages of development including mature eggs (data not shown).

**Figure 3.**
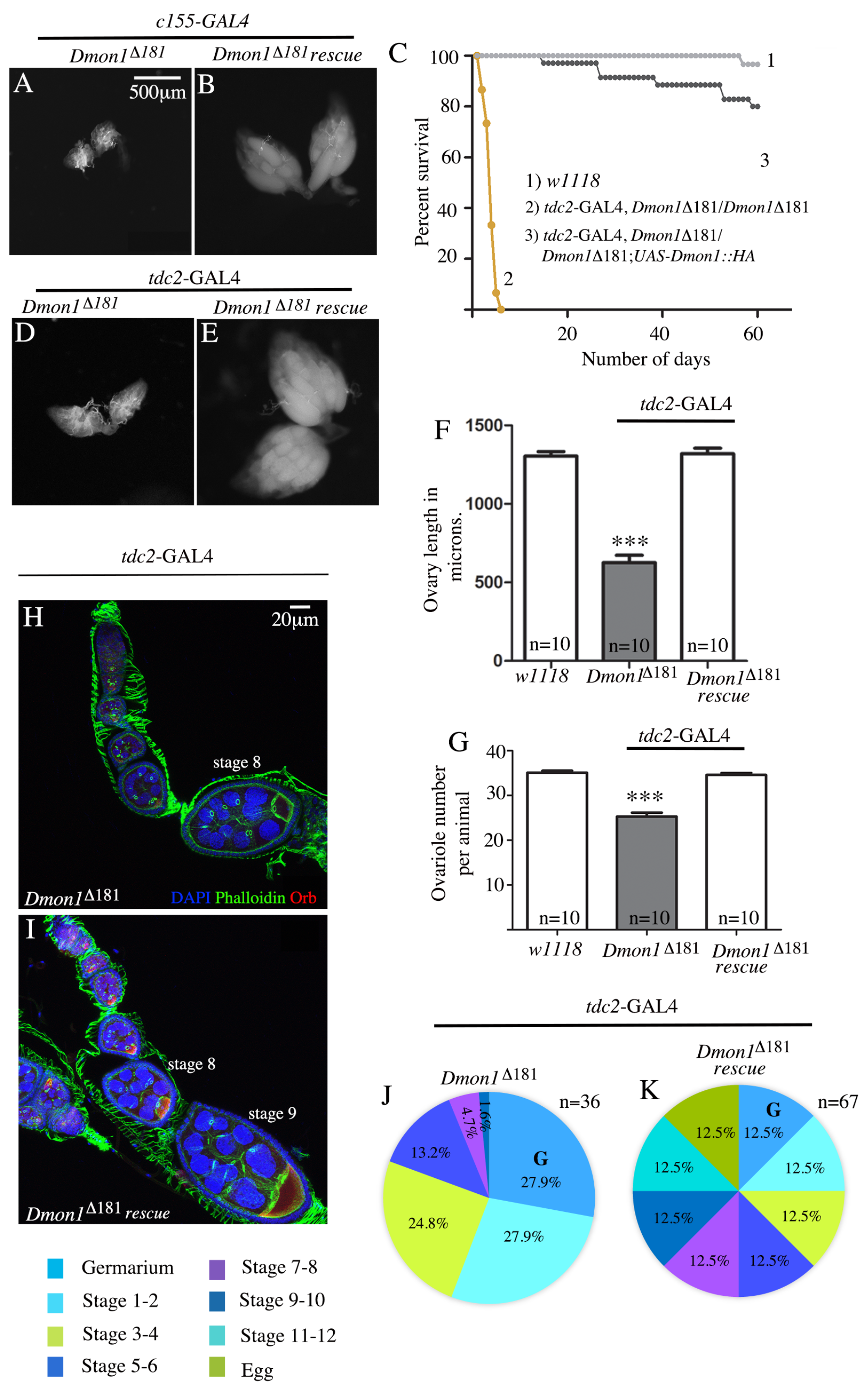
Expression of *Dmon1* in OPNs is sufficient to rescue lethality and ovarian defects in *Dmon1 ^Δ181^*. **(A, B)** Pan-neuronal expression of *DMon1:HA* in *Dmon1^Δ181^* mutants (B, c155-GAL4; *Dmon1^Δ181^/Dmon1^Δ181^; UAS-Mon1HA*) rescues the ‘small ovary’ phenotype. **(C, D)** Expression of *DMon1:HA* in *Dmon1^Δ181^* mutants using *tdc2*-GAL4 (B, *tdc2*-GAL4, *Dmon1^Δ181^/Dmon1^Δ181^; UAS-Mon1HA*) rescues the ‘small ovary’ phenotype. **(E)** Lethality of *Dmon1^Δ181^* mutants is rescued by expression of the gene in the *tdc2* domain (*tdc2*-GAL4, *Dmon1^Δ181^/Dmon1^Δ181^; UAS-Mon1HA)*). **(F, G)** Expression of *Dmon1:HA in* OPNs rescues the defect in ovary size (F) and ovariole number observed in *Dmon1^Δ181^* animals. **(H-K)** Expression of *Dmon1:HA* in OPNs (*tdc2*-GAL4, *Dmon1^Δ181^/Dmon1^Δ181^; UAS-Mon1HA*) rescues the ‘stalling’ defect in *Dmon1^Δ181^* ovaries. Note the presence of late stage egg chambers in a ‘rescued’ ovariole (I). Orb expression is also restored in these egg chambers (Compare panel H with I). (J, K) The distribution of egg chambers in ovaries from *tdc2*-GAL4, *Dmon1^Δ181^/ Dmon1^Δ181^* (J) and *tdc2*-GAL4, *Dmon1^Δ181^/ Dmon1^Δ181^; UAS-Mon1HA* animals (K). Note the dramatic rescue of egg chamber distribution (panel K) which resembles that of wild type (Fig. 1L).

Next, we tested if expression of *Dmon1*, in the OPNs alone, using *tdc2*-GAL4 can rescue lethality and the sterility defect in *Dmon1^Δ181^*. Surprisingly, under these conditions a complete rescue of lethality was observed. While homozygous mutant flies fail to survive beyond 7 days, the ‘rescued’ adults were able to survive well beyond 30 days (Fig. 3C). Moreover ovaries from such ‘rescued’ animals were morphologically similar to wild type (Fig. 3E). The length of these ovaries, used as an indicator of size, was found to be comparable to wild type control females: 1305 ±28.04 *μm* (*w^Δ118^*) *vs* 627.5±45.84 *μm* (*tdc2*-GAL4,*Dmon1^Δ181^/Dmon1^Δ181^), 1319±35.77μm (Dmon1^Δ181^, *tdc2*-GAL4/Dmon1^Δ181^; UAS-Mon1-HA)* (Fig. 3F). Ovariole number was also rescued in these animals (35.10±0.43 (n=10, w^1118^) vs 25.30±0.84 (n=10, *Dmon1^Δ181^*, *tdc2*-GAL4 / *Dmon1^Δ181^) versus 34.60±0.43 (n=10, Dmon1^Δ181^, *tdc2*-GAL4 / Dmon1^Δ181^; UAS-Mon1-HA*) (Fig. 3G). Presence of late stage egg chambers including eggs were seen in the rescued ovarioles and Orb levels appeared to be restored (Compare Fig. 3H and 3I). We looked at the distribution of egg chambers and as seen in case of rescue with *c155-Gal4*, these animals (*Dmon1^Δ181^*, *tdc2*-GAL4, */ Dmon1^Δ181^; UAS-Mon1-HA)* showed complete rescue of the ‘stalling’ defect (Compare Fig. 3J and 3K). Degeneration of egg chambers was also rescued (data not shown). Thus, expression of *Dmon1* in *tdc2*-Gal4 domain was sufficient to rescue all aspects of the ovarian phenotype observed in *Dmon1^Δ181^* mutants.

The ability to rescue the *Dmon1* dependent sterility by expression of *Dmon1* in the OPNs revealed an unanticipated yet close functional connection between *Dmon1* activity and the particular neuronal subtype. Importantly, expression of *Dmon1* in the OPNs allowed oogenesis to progress beyond the pre-vitellogenic stages. In the following section’s we present our efforts to elucidate the possible mechanistic basis underlying the influence of OPNs during oogenesis.

### Expression of Drosophila insulin like peptides (Dilps) is downregulated in Dmon1 mutants

Arrested oogenesis is a hallmark of insulin pathway mutants. In particular, the role of insulin signaling during ovarian development in *Drosophila melanogaster* is well established (16). Mutations in *chico*, the substrate for insulin receptor, also cause sterility and prevent transition of the egg chambers from pre-vitellogenic to the vitellogenic phase (14) resulting in smaller sized ovaries.

It has been previously demonstrated that OPNs can influence IPCs and, thereby can modulate insulin production and downstream signaling (18, 29). Given the broad similarity in the ovarian phenotypes between *Dmon1^Δ181^* and Insulin pathway mutants, we wondered if *Dmon1* activity may target one or more of insulin like peptides. We therefore decided to test if endogenous insulin levels and/or signaling are compromised in *Dmon1* mutants. There are eight insulin like peptides in *Drosophila*. As a first test of this model, we quantitated the expression levels of these *dilps* in total RNA samples made from mutant female adults using quantitative RT-PCR. While no significant change was observed in the expression of *dilp2* and *dilp7*, a significant decrease in expression was seen the case of *dilps* 3,4,5 and 6 and an increase in the case of 8 as opposed to wild type ovarian RNA samples (Fig. 4A).

**Figure 4.**
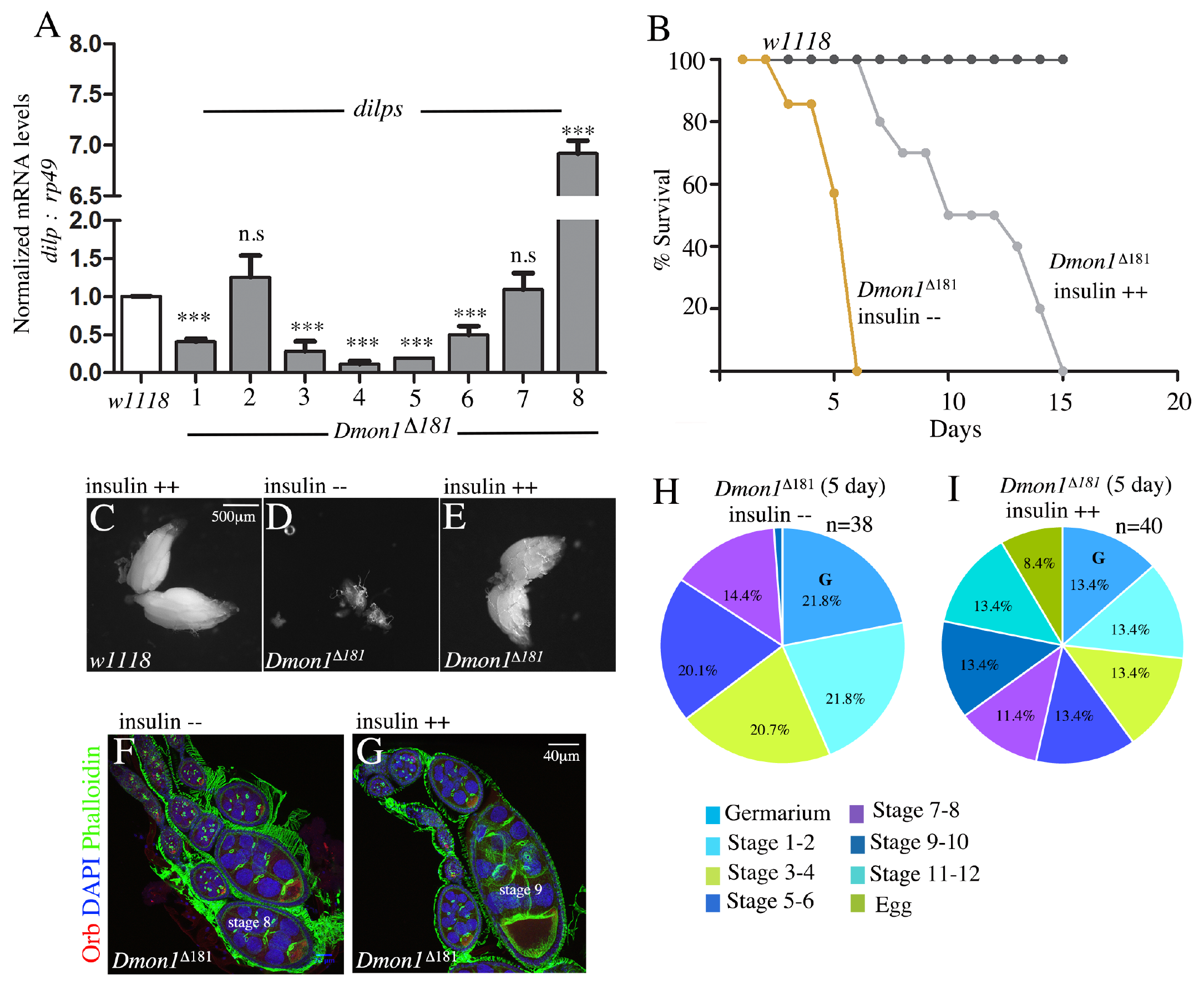
Feeding insulin to *Dmon1^Δ181^* mutants supresses the ovary defects. **(A)** Estimation of *dilp* mRNA levels from 2-3 day old adult *Dmon1^Δ181^* animals normalized to wildtype using quantitative real time PCR. *dlips* 1,3,4,5,6, and 8 show significant changes in transcript levels in the *Dmon1* mutants. **(B)** Life span of *Dmon1^Δ181^* mutants maintained on normal (*Dmon1^Δ181^* insulin-ve) and 5ug/ml insulin containing medium (*Dmon1^Δ181^* insulin +ve). The half-life of survival is doubled in animals fed on insulin containing medium. **(C-E)** Shown are 5 day old ovaries from wildtype (C, insulin positive) and *Dmon1^Δ181^* mutants kept on normal (D) and insulin containing medium (E). Note the restoration of ovary size in mutants upon insulin feeding (E). (F, G) Ovarioles from *Dmon1^Δ181^* (Ίnsulin-ve) (F) and *Dmon1^Δ181^* (insulin+ve) mutant (G) stained with Orb (red), DAPI (blue) and Phalloidin (green). Note the presence of late stage egg chambers in (G). **(H, I)** Pie-charts showing the proportion of the different stages of egg chambers in ovaries from *Dmon1^Δ181^* (insulin negative, H) and *Dmon1^Δ181^* (insulin positive, I) animals. Note the appearance of late stage egg chambers in proportions comparable to wildtype.

### Insulin rich diet can suppress mon1 dependent female sterility

Given the significant reduction in *dilp levels* in *Dmon1* mutants, we wondered if feeding insulin-rich diet to homozygous mutants will ameliorate the oogenesis defects. When freshly eclosed *Dmon1^Δ181^* mutants were transferred to medium containing 5ug/ml of insulin (see Methods), we observed a significant improvement in the lifespan of these animals. While most homozygous *Dmon1^Δ181^* mutants die between day 5 and day 7, the mutants maintained on insulin containing medium were able to survive till day 15 (Fig. 4B). We examined ovaries of 5-day old insulin fed mutants with age-matched mutants that were maintained on normal medium. In contrast to mutant ovaries (Fig. 4D), the ovaries from the insulin fed flies appear to be nearly normal (Fig. 4E) and comparable to ovaries from wild type animals fed on insulin (Fig. 4C). Importantly, ovarioles from insulin fed flies displayed late stage egg chambers (Fig. 4F, G). This was evident upon plotting the distribution of the different stages: ovarioles from homozygous 5-day old *Dmon1^Δ181^* mutant failed to develop egg chambers beyond stage 7 or 8; the ovarioles from insulin fed flies showed presence of all stages (Fig. 4I) similar to a wild type ovary sample (See Fig. 2). To confirm that the effects of insulin are not due to increase in the amount of protein in the medium, we carried out experiments in which mutant flies were separately maintained either on yeast rich medium or on medium containing equivalent amount of FLAG peptide (Sigma, F3290). In both cases, the ovaries appeared similar to those from mutant animals maintained on normal medium. This suggests that the suppression of the ovary phenotype in insulin fed flies is not due to a nutrient or amino-acid ‘rich’ diet but likely because of insulin activity.

### *DLIP5* levels are considerably reduced in Dmon1 mutant IPCs

We find the transcript levels of a number of *dilps* are substantially reduced in *Dmon1* mutants (Fig. 4A). Thus, suppression of *Dmon1* phenotype by feeding insulin-rich diet is an indication of its ability to influence the level of one (or more) DILPs produced by the IPCs. Alternatively, the improved morphology and progression through vitellogeneis upon insulin feeding could be simply a result of ‘insulin dependent bypass’ mechanism. It was thus important to examine if *Dmon1* is specifically necessary to maintain DILP levels in the IPCs. IPCs synthesize DILP2, 3 and 5 but only *dilp3* and *5* RNA levels are reduced in *Dmon1* mutants. Since *dilp5* has been shown to influence oogenesis (14, 30), we decided to examine the level of DILP5 in IPCs by staining the adult brains using anti-DILP5 antibodies. As can be seen (Fig. 5B) there is a substantial decrease in DILP5 in *Dmon1^Δ181^* IPCs as compared to the heterozygous mutant (Fig. 5A). We quantitated the intensity of DILP5 staining in the IPCs and compared the levels to that in heterozygous mutant as well as wildtype. In both cases, we observed a 60% decrease in DILP5 intensity. Interestingly, a similar decrease in DILP5 level is also observed in IPCs when *Dmon1* activity is compromised only in OPNs (Compare Fig. 5D and 5E). A 40% reduction in the intensity of DILP5 staining was observed in these animals (Fig. 5F). *dilp2* mRNA levels are not significantly altered in *Dmon^Δ181^* mutants (Fig. 4A).

**Figure 5.**
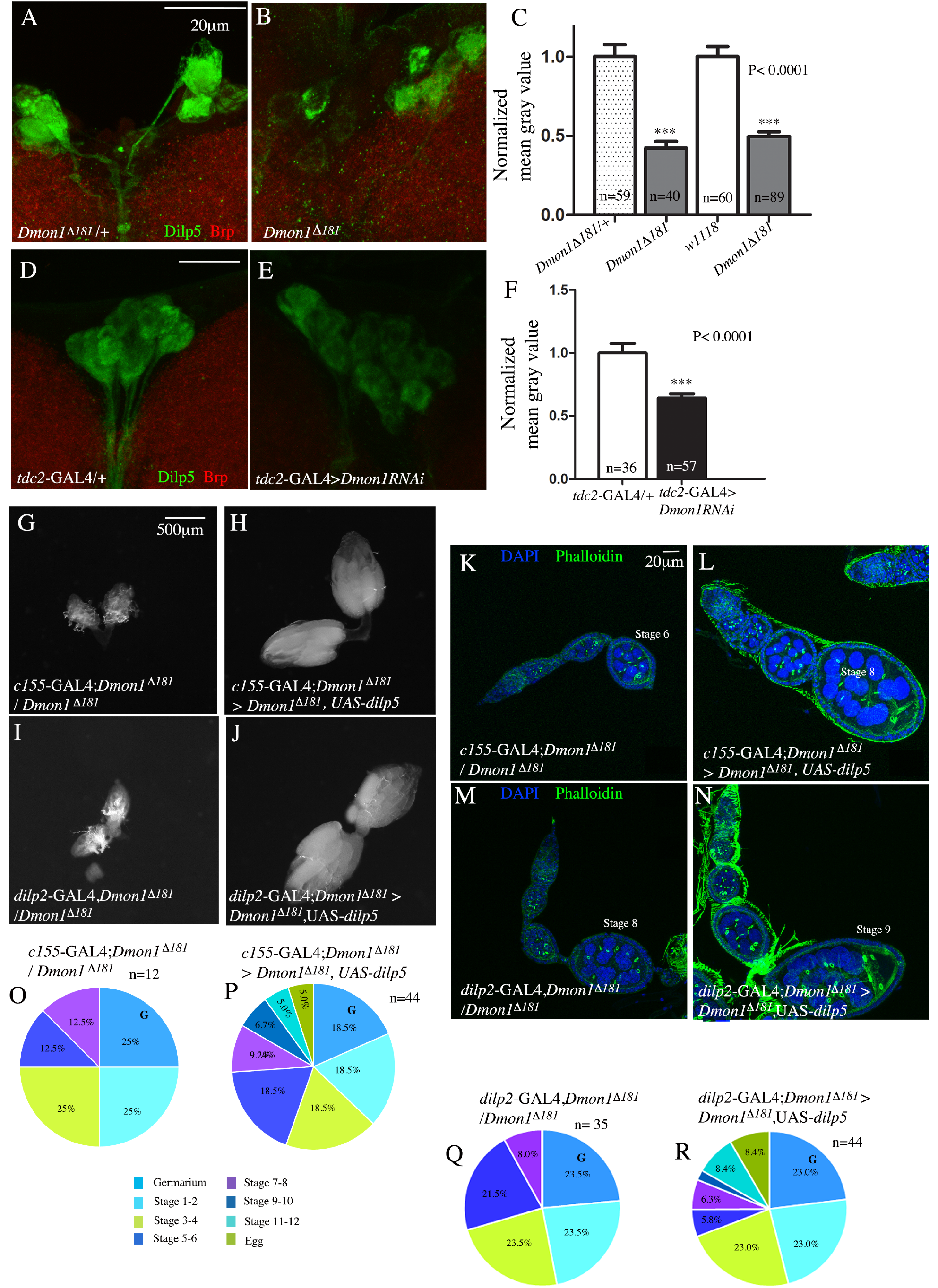
*Dmon1* in OPNs regulates *dilp5* which can suppress the ovary defects in *Dmon1^Δ181^*. **(A-C)** Adult brains of *Dmon1^Δ181^*/+ (A) and *Dmon1^Δ181^* mutants (B) stained with anti-Dilp5 (green) and anti-Brp (red). Intensity of Dilp5 staining is reduced by nearly 50% in the mutants (C). **(D-F)** Adult brains of *tdc2*-GAL4/+ (D) and *tdc2*-GAL4>*UAS-Mon1RNAi* (E) stained with anti-Dilp5 (green) and anti-Brp (red). Intensity of Dilp5 staining in the IPCs is reduced in upon downregulation of *Dmon1* in OPNs (Compare E to D). The decrease in intensity is approximately 40% (F). **(G-H)** Pan-neuronal expression of *dilp5* (*c*155-GAL4; *Dmon1^Δ181^/Dmon1^Δ181^, UAS-dilp5*) rescues ovary size in *Dmon1^Δ181^* mutants (Compare H to G). **(I-J)** Expression of *dilp5* in IPCs using *dilp2*-GAL4 (*Dmon1^Δ181^,dilp2*-GAL4/ *Dmon1^Δ181^*,*UAS-dilp5*) rescues ovary size in *Dmon1* mutants (Compare J to I). **(K-M)** Representative ovarioles from *c*155-GAL4; *Dmon1^Δ181^/; Dmon1^Δ181^, UAS-dilp5* (L) and *dilp2*-GAL4, *Dmon1^Δ181^/; Dmon1^Δ181^, UAS-dilp5* (N) animals stained with DAPI and phalloidin. Note the presence of late stage egg chambers, which are absent in their respective controls (K, M). **(O-R)** Pie-charts showing the distribution of different stages of egg chambers in *Dmon1^Δ181^* mutants rescued with *UAS-dilp5* either pan-neuronally (P) or through expression in the IPCs (R). Late stage egg chambers are seen in both cases compared to their respective controls (O and Q).

### Overexpression of dilp5 in IPCs can alleviate loss of Dmon1

We thus sought to assess if the reduction in DILP5 levels in *Dmon1* IPCs is an important factor that contributes to its ability to regulate egg chamber maturation. If so, neuronal expression of DILP5 in the IPCs should be able to mitigate the ovarian phenotype due to loss of *Dmon1*. This was indeed the case. Expression of *UAS-dilp5* either using a pan-neuronal driver (*C155-Gal4;* compare Fig. 5G and 5H) or in the IPCs alone (*dilp2-Gal4;* compare Fig. 5I and 5J) was able to ameliorate the phenotypes of *Dmon1^Δ181^* considerably. The animals showed improved viability and a suppression of the ovarian phenotype including sterility. In both cases, ‘rescued’ ovaries showed ovariole numbers comparable to wild type and presence of late stage egg chambers (Compare Fig. 5K and 5L; also Fig. 5M and 5N). A quantitation of the proportion of the different stages of egg chambers showed that while both ‘driver control’ mutants (*c155-GAL4; Dmon1^Δ181^/ Dmon1^Δ181^* and *tdc2*-GAL4, *Dmon1^Δ181^/ Dmon1^Δ181^*) exhibit arrested egg chambers at stage 7-8 (Fig. 5O and 5Q respectively), expression of *dilp5*, relieves this stalling resulting in the development of late stage egg chambers (Fig. 5P and 5R respectively). Interestingly, unlike the rescue with *UAS-Mon1::HA*, the proportion of early egg chambers does not appear to change in *tdc2*-GAL4, *Dmon1^Δ181^/ Dmon1^Δ181^, UAS-dilp5* animals suggesting a developmental delay either due to insufficient levels of DILP5 or requirement of additional Dilps. Confirming the IPC specific role of DILP5, over expression of *dilp5* in OPNs was unable to suppress the loss of viability and ovary phenotypes in the mutants.

### Can other DILPs substitute for DILP5 loss in Dmon1 mutants?

IPCs are known to synthesize DILP2, 3 and 5. Previous studies have indicated existence of partial redundancy among different DILPs (31, 32). To examine if DILP2 expression can achieve an analogous rescue of *Dmon1* phenotype, we expressed *UAS-dilp2* in the IPCs as well as using a pan-neuronal driver. Expression of *dilp2* in *Dmon1^Δ181^* mutants was however unable to alleviate the mutant phenotype. While we have not been able to test *dilp3* in a similar manner, our data show that *Dmon1* functions in the OPNs to regulate the levels of DILP5 in the IPCs (Fig. S1). Furthermore, *dilp5* is one of the important targets of *Dmon1* as oogenesis defects induced by loss of *Dmon1* activity can be substantially suppressed by expressing *dilp5* in IPCs.

## Discussion

DILPs in conjunction with the insulin receptor have been shown to be involved in a variety of conserved biological processes that ultimately impact growth advantage, developmental profile, reproductive potential and aging dependent decline (14, 33, 34). Recent data have also suggested that insulin can affect behavioral traits via nervous system function and physiology (16, 35).

In *Drosophila melanogaster*, differential expression of the DILPs underlies diverse functions in different tissue contexts. A number of transcriptional mechanisms are thought to be involved in conferring tissue specific expression of individual *dilps* and hence the pathways responsible for activation of IPCs as well as modes of transport and reception of DILPs are under rigorous investigation.

Emphasizing the importance of long distance systemic communication involved in organogenesis, elegant studies by Rajan and Perrimon demonstrated that fat body cells synthesize JAK/STAT ligand, Unpaired2 (Upd2) which, in turn, communicates with the IPCs to induce secretion of *dilps* (36). Subsequent analysis from Leopold lab elucidated IPC specific requirement of a G protein coupled receptor Methuselah (Mth) that acts in conjunction with fat body cell derived ligand Stunted (Sun) to regulate proper secretion of DILPs (37). Here we have focused our attention on an endocytic regulator, DMon1 that appears to participate in a similar ‘remote control’ type of mechanism. Interestingly however, DMon1 does not seem to function in the IPCs but rather is needed in an adjacent neuronal population i.e. OPNs, to ultimately achieve physiologically relevant insulin levels crucial for systemic communication underlying proper organogenesis (Fig.6).

*Dmon1* mutants display female sterility as the mutant females have small ovaries with diminished number of ovarioles. The individual mutant egg chambers arrested at the previtellogenic stages and mutant ovaries are devoid of any mature eggs. Interestingly, *Dmon1* activity appears to be required not in the ovaries, but in the nervous system. Supporting such a non-cell autonomous control by DMon1, compromising *Dmon1* levels specifically in the OPNs is sufficient to induce the ‘small ovary’ phenotype. Importantly, expression of *Dmon1* pan-neuronally or in the OPN neurons alone, is sufficient to rescue sterility and associated phenotypes induced by the loss of *Dmon1*.

Since several of the ovarian phenotypes including sterility are reminiscent of mutants in the insulin signaling pathway, we examined the possible connection between *Dmon1* and Insulin signaling. Two related findings are of interest in this regard. Firstly, levels of a number of *dilps (1, 3, 5* and 6) are significantly reduced in *Dmon1^Δ181^* mutants. Importantly, feeding insulin rich diet to homozygous *Dmon1^Δ181^* females ameliorates the ovarian phenotypes induced by the loss of *Dmon1*.

The observation that expression of *Dmon1* using a pan-neuronal driver is sufficient to rescue both lethality and the sterility phenotype strongly supports a neural basis for the *Dmon1* dependent defects. Our data also argue that OPNs are likely the functionally critical subset in this regard. OPNs are known to modulate many behavioral traits in *Drosophila* (18, 29). Taken together our data suggest that these neurons likely control IPCs to ultimately modulate DILP production and/or transmission. In this regard, it is noteworthy that the terminals of some of the OPNs lie in close proximity to the IPCs which secrete DILPs 2, 3 and 5 (18). Moreover, IPCs are also known to express OAMB OPN receptors on their surface opening up an exciting possibility that IPCs can directly respond to octopamine activity and/or levels. The feeding rescue with insulin rich diet further argues that *Dmon1* in OPNs may be necessary for the synthesis and/or secretion of DILPs.

Our data suggests that transcript levels of *dilp3* and 5 (but not *dilp2*), are attenuated in *Dmon1* mutants. Such a reduction suggests that loss of DMon1 could affect the transcription of these genes (*dilp3 and* 5). Whether the circulating levels of the respective DILPs are also affected in a consistent manner still remains to be determined. Our genetic analysis has indicated that expression of DILP2 in *dilp2* expressing neurons is not sufficient to rescue the lethality and ovarian phenotype in *Dmon1^Δ181^* mutants (data not shown). It is thus possible that *Dmon1* activity targets specific members of the *dilp* family. Nevertheless, altogether our results argue that *Dmon1* is genetically upstream of insulin signaling, likely functions in a subset of OPNs and future experiments will focus on elucidating the mechanistic underpinnings of this regulatory relationship.

It is remarkable that in the neuromuscular junction context, *Dmon1* appears to be secreted by the presynaptic terminals (23). This observation suggests that it is competent to function in a non-cell autonomous manner. As in the case of neuromuscular junctions, it is possible that DMon1 engineers the expression of OAMB thereby indirectly influencing the activation of IPCs. Alternatively, DMon1 may facilitate the release of as yet unknown factors that influence expression of dilps/DILPs. These possibilities need to be explored by directly testing the levels and/or activity of OAMB receptors and *dilps* in the IPCs.

Secretion of DILPs is thought to be regulated by ecdysone and juvenile hormone. We therefore decided to test if DMon1 exerts its influence on oogenesis via these pathways. Arguing against the possibility, however, feeding *Dmon1^Δ181^* mutant flies with 20-OH ecdysone did not rescue the ovarian phenotype to any appreciable extent. By contrast, feeding excess levels of insulin effectively rescued the *Dmon1^Δ181^* dependent ovarian phenotype. Furthermore, the extent of rescue can be correlated with the amount of insulin intake and the length of feeding time. For instance, we find that mutant flies fed on insulin containing medium for 10 days, show a near complete rescue of the ovarian phenotypes including rescue of the ovariole number and presence of mature eggs. A similar rescue is seen at day 5 although these ovaries tend to have a few ‘mutant-like’ small ovarioles. Taken together these observations strongly suggest that defects induced by the loss of DMon1 are, at least in part, due to impaired insulin signaling.

In summary, our data have revealed a novel regulatory loop (Suppl. Fig. S1) involving a specific subset of neurons and insulin metabolism which, in turn results into phenotypic abnormalities at the organismal level. While the precise identity of the individual effectors is unclear at present, these data clearly demonstrate how systemic level communication between individual cells, tissues, and even organs can participate in, and contribute to, organismal homeostasis.

## Experimental methods

### Fly husbandry and strains used

All stocks were reared on standard corn meal agar medium. *Dmon1^Δ181^* and *Dmon1^Δ129^* were generated through excision of pUAST-Rab21:: YFP insertion using standard genetic methods (23). *Df(2L)9062, c*155-GAL4, *tdc*\2-GAL4 (#9313), *htl*-GAL4 ^GMR93H07^ (#40669) are from Bloomington Stock Centre, Indiana, USA; nos-GAL4 (R.Rikhy, IISER-Pune); *dilp2-GAL4* (Rulifson et al., 2002); *Dmon1* RNAi line (GD7852) is from the VDRC stock centre; *UAS-mon1:: HA* (kind gift from T.Klein) All crosses were carried out at 25 degrees.

### Immunostaining and Imaging

For whole ovary immunostaining and imaging standard protocols were used (see Suppl. text for a detailed description). The concentrations of the different dyes and antibodies were as follows: DAPI (Invitrogen Molecular Probes, 1:500); Phalloidin 488 (Invitrogen, Molecular Probes, 1:100); Anti-Orb (DSHB, 1:20); Alexa flour secondary antibodies (Invitrogen Molecular Probes, 1:1000). Imaging was done using the Leica SP8 confocal microscope system with a 40X oil objective (1.3 N.A.) Immunostaining of adult brains was carried out as described in (38). Anti-DILP5 (kind gift from P. Leopold, France) was used at 1:800. Image processing and intensity measurements were done using Image J software (NIH). GraphPad Prism was used for statistical analyses. Figures were assembled using Adobe Photoshop CS4.

### Quantitative RT-PCR

Estimation of *orb* levels was carried out using quantitative PCR on cDNA synthesized from mRNA isolated from adult wildtype (w1118) and *Dmon1* mutant ovaries. The ovaries were dissected in chilled 1XPBS. RNA isolation was carried out using Trizol reagent as per the manufacturer’s instructions. For estimation of *dilps*, mRNA was isolated from 1-2 day old adult female flies. In each case, 1 μg of RNA was used for the reverse transcription reaction. The sequence of the primers used for quantitative PCR are listed as Suppl. Text.

### Insulin Feeding experiments

Regular corn meal medium containing 5μg/ml of bovine insulin (Sigma Aldrich, I6634). Newly eclosed wildtype and *Dmon1^Δ181^* mutant females were transferred to insulin containing medium. Ovaries were examined in 5 day old animals. For measurement of life span, the animals were maintained on insulin medium for the entire duration of the experiment.

## Acknowledgements

We thank the Bloomington Drosophila Stock Centre and the Vienna Drosophila RNAi centre for fly stocks; Pierre Leopold for DILP antibodies; ARI and IISER, Pune, for use of the Confocal microscopy facility; NCBS, Bangalore, and C-CAMP, Bangalore for stocks; Gaiti Hasan, and LS Shashidhara for helpful discussions; This work was supported by funds from Department of Biotechnology, Govt. of India (GOI) and intramural funds from ARI, Pune to AR; intramural funds from IISER, Pune, to GR. G.D is supported by a grant from NIH (GM110015).

## Author Contributions

AR conceived the project and designed the experiments; AR and ND performed the experiments; AR, ND, GR and GD analyzed the data; AR, GD and GR wrote the manuscript.

---

**Figure S1.**
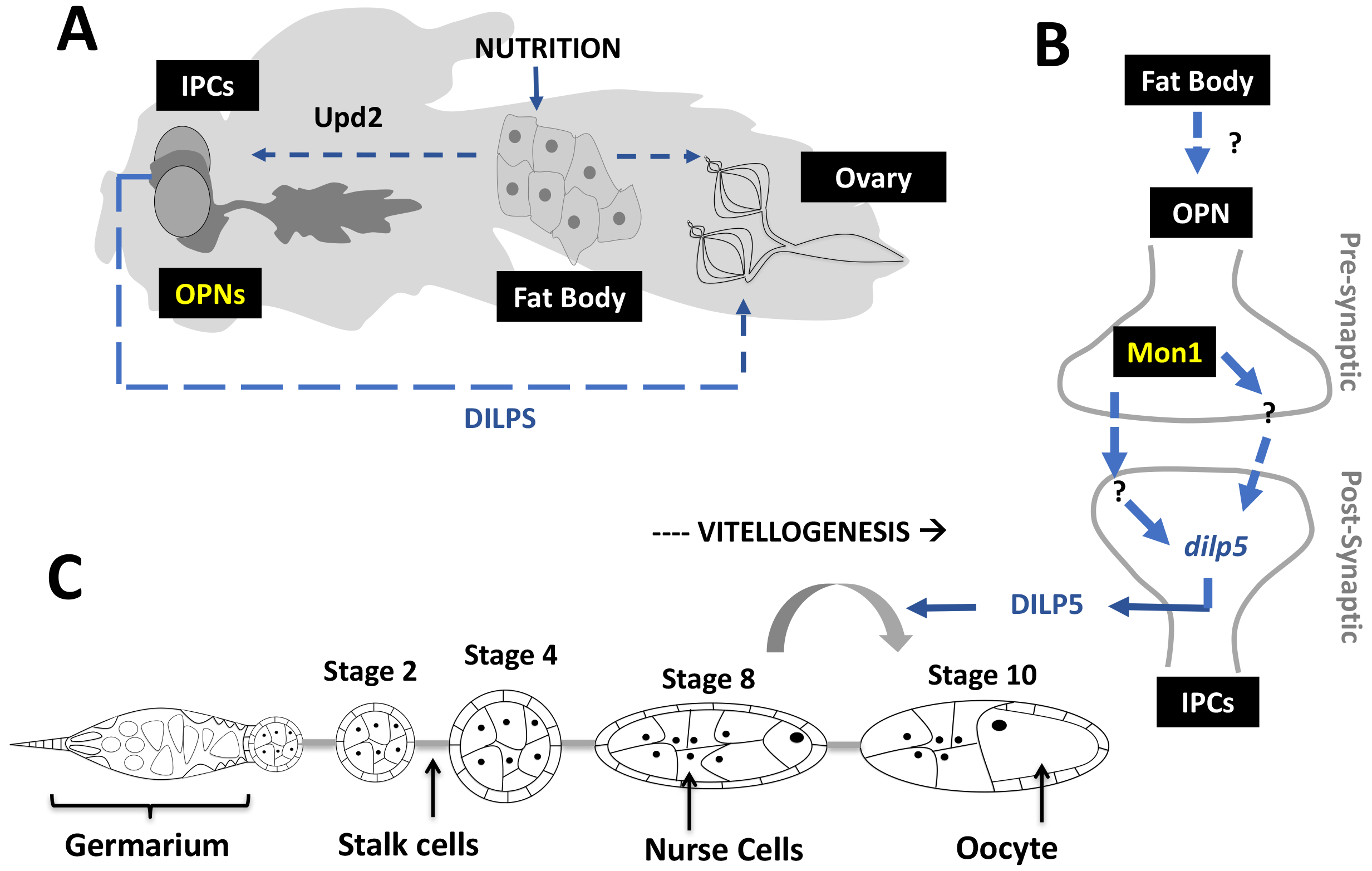
*Mon1* activity in the OPNs modulates DIP5 levels to influence vitellogenesis and ovarian maturation. **(A)** In the adult fly, Insulin like peptides from IPCs regulate development of egg chambers. The expression/secretion of DILPs from the IPCs in turn depends on nutritional cues sensed by the Fat Body and communicated to the IPCs via Upd2, a JAK-STAT ligand. Upregulation of Upd2 relieves the inhibitory action of GABA neurons on the IPCs, which in turn induces the IPCs to release DILPs. OPNs have not been implicated in ovarian development. **(B)** Our data suggest that OPNs, signal to and regulate *dlip5* levels in the IPCs. This activity depends on *Dmon1* in the OPNs. The precise identity of the signal that engineers the communication between OPNs and IPCs is currently unknown. Mon1 is thus an important player in the brain-gonad axis, critical for proper ovarian development, which in turn is dependent on response to nutritional as well as hormonal cues. **(C)** DMon1 in OPNs acts as a remote-control mechanism that modulates *dilp5* expression to achieve proper oogenesis, specifically acting to regulate transition of egg chambers from previtellogenic to vitellogenic stages. Importantly, this regulation appears to function under normal feeding conditions i.e. in the absence of starvation.

